# Evaluation in mice of cell-free produced CT584 as a Chlamydia vaccine antigen

**DOI:** 10.1101/2024.06.04.597210

**Authors:** Steven Hoang-Phou, Sukumar Pal, Anatoli Slepenkin, Abisola Abisoye-Ogunniyun, Yuliang Zhang, Sean F. Gilmore, Megan Shelby, Feliza Bourguet, Mariam Mohagheghi, Aleksandr Noy, Amy Rasley, Luis M. de la Maza, Matthew A. Coleman

## Abstract

*Chlamydia trachomatis* is the most prevalent bacterial sexually transmitted pathogen worldwide. Since chlamydial infection is largely asymptomatic with the potential for serious complications, a preventative vaccine is likely the most viable long-term answer to this public health threat. Cell-free protein synthesis (CFPS) utilizes the cellular protein manufacturing machinery decoupled from the requirement for maintaining cellular viability, offering the potential for flexible, rapid, and de-centralized production of recombinant protein vaccine antigens. Here, we use CFPS to produce the putative chlamydial type three secretion system (T3SS) needle-tip protein, CT584, for use as a vaccine antigen in mouse models. High-speed atomic force microscopy (HS-AFM) imaging and computer simulations confirm that CFPS-produced CT584 retains a native-like structure prior to immunization. Female mice were primed with CT584 adjuvanted with CpG-1826 intranasally (i.n.) or CpG-1826 + Montanide ISA 720 intramuscularly (i.m.), followed four-weeks later by an i.m. boost before respiratory challenge with 10^4^ inclusion forming units (IFU) of *Chlamydia muridarum*. Immunization with CT584 generated robust antibody responses but weak cell mediated immunity and failed to protect against i.n. challenge as demonstrated by body weight loss, increased lungs’ weights and the presence of high numbers of IFUs in the lungs. While CT584 alone may not be the ideal vaccine candidate, the speed and flexibility with which CFPS can be used to produce other potential chlamydial antigens makes it an attractive technique for antigen production.

## Introduction

*Chlamydia trachomatis* (*Ct*) is the most common bacterial sexually transmitted pathogen worldwide and a large public health threat^1^. Untreated infections, particularly in women, can develop serious sequelae including pelvic inflammatory disease and infertility^2,3^. While the rates of infection have trended downward in the 2000’s, it has been reversed over the last decade^4^. Alarmingly, the SARS-CoV-2 pandemic resulted in a scarcity of chlamydia testing and tracking, with predicted *Ct* case rates exceeding actual measured rates, leading to the possibility of missed diagnoses^5^. Since *Ct* infections are often asymptomatic, community surveillance is key to understanding prevalence, and it is clear that a vaccine against *Ct* would be the most effective approach to limit infection, spread, and protect public health.

Many different chlamydial antigens have been tried in vaccine development and it remains an area of active investigation. Whole-cell fixed or attenuated chlamydial elementary bodies (EBs) offer some of the best protection from subsequent infections in mouse models^6^, but ambiguity in the safety, generated responses, and scalability in the context of mass human vaccinations remain an important barrier^7^. Recombinant protein antigens have seen some success in animal vaccine trials, largely focused on the major outer membrane protein (MOMP) which comprises ∼60% of the mass of the chlamydial outer membrane^8^. While formulations using variations of this antigen have shown the most promise, other protein antigens present on the cell surface have been tested with varying results (Reviewed in ^9^).

One promising candidate is the chlamydial type III secretion system (T3SS) “injectosome” protein complex. The T3SS is responsible for the translocation of effectors from the chlamydial cytosol into the host cell cytoplasm, promoting chlamydial virulence^10^. It is comprised of over 10 different proteins, partially protruding from the chlamydial cell surface, presenting a few potential antigenic targets. Importantly, the T3SS is present at all stages of the chlamydial life cycle^11^, and inhibition of the T3SS has been demonstrated to reduce chlamydial infection in cells^12–14^. Recently, a recombinant protein antigen consisting of a CopB, CopD, and CT584 fusion has been shown to confer partial protection in both *Chlamydia muridarum* (*Cm*) and *Ct* challenge studies in mouse models^15,16^. The proteins CopB and CopD are thought to form the terminal translocon pore while CT584 has been suggested to be the needle tip protein of the T3SS^17–19^.

CT584, as a putative needle tip protein in the chlamydial T3SS, makes an excellent target for potential vaccine formulations. Targeting and inhibition of other needle tip proteins, such as LcrV in *Yersinia pestis*, prevents secretion of translocon and effector proteins, precluding apoptosis of infected cells^20,21^. CT584 from *Ct* serovar D is highly conserved (97.27% identical) at the amino acid sequence level with its homolog TC873 in *Cm* (Supplementary Figure S1). Since *Cm* infection mouse models produce similar infection and long-term sequelae to *Ct* infection in humans^22,23^ and vaccine formulations containing *Ct* CT584 have shown promising results in both *Cm* and *Ct* mouse models^15,16^, we hypothesized using CT584 alone as a vaccine antigen in *Cm* mouse models may induce protective adaptive immune responses that limit infection and pathology and could potentially translate well to the clinic.

Depending on the protein being tested, antigen production for immunization studies can be challenging. Cell-free protein synthesis (CFPS) offers a rapid and flexible way to express and test potential protein antigens. Coupled transcription-translation CFPS systems utilize cell lysates containing an RNA polymerase, ribosomal complexes, amino acids, nucleotides, and typically an ATP regeneration system to maintain protein synthesis, while a plasmid DNA template is added to direct mRNA synthesis and protein production^24^. *Escherichia coli*-based lysates are currently the best characterized and highest performing system for CFPS applications, yielding >2mg protein per mL of reaction in less than 16 hours (Reviewed in ^25^). CFPS is also scalable, offering the ability to rapidly switch gears and produce other proteins – only the DNA template needs to be changed. CFPS methods are gaining popularity due to their ease of use, scalability, and rapid production, and the first CFPS-produced therapeutics^26,27^ and vaccines^28^ are entering clinical trials.

Here, we applied *E. coli*-based CFPS to rapidly produce and then characterize recombinant CT584 protein based on the *Ct* serovar D amino acid sequence as a potential vaccine antigen. We produced >1 mg of pure, native-like, protein per mL CFPS reaction for use in immunization and testing in mouse models of chlamydial infection. We show that robust IgG antibody responses to CT584 immunization are generated using different routes of immunization and antigen/adjuvant formulations, however, the formulations of CT584 were not protective against respiratory *Cm* challenges. Furthermore, we did not detect CT584 protein in chlamydial EB using EB immune sera while CT584 immune sera could detect it, suggesting that CT584 is naturally a weak immunogen.

## Results

### High yields of CT584 can be produced using cell-free protein synthesis

Optimal production of poly-histidine tagged *Ct* CT584 (hereafter CT584) through CFPS requires a highly productive cell-free lysate, reaction buffer and energy regeneration solution, and plasmid containing the codon optimized *ct584* DNA template (Figure 1A). Codon optimization introduces silent mutations into the DNA coding sequence of the template to better match tRNA frequencies and translation speeds encountered in the host cell (*E. coli*) lysate to improve translation efficiency^29^. Codon optimization of the *Ct* serovar D derived CT584 sequence resulted in the replacement of 25.14% of its DNA bases (Figure 1B, Supplementary Figure S2). Nascent CT584 can be labeled through inclusion of a BODIPY dye-labeled tRNA-Lys in the cell-free reaction, which incorporates labeled lysine residues into actively made proteins. Specific production and high yields of CT584 can be seen through BODIPY fluorescence and SYPRO staining for total protein on SDS-PAGE gels (Figure 1C).

**Figure 1.**
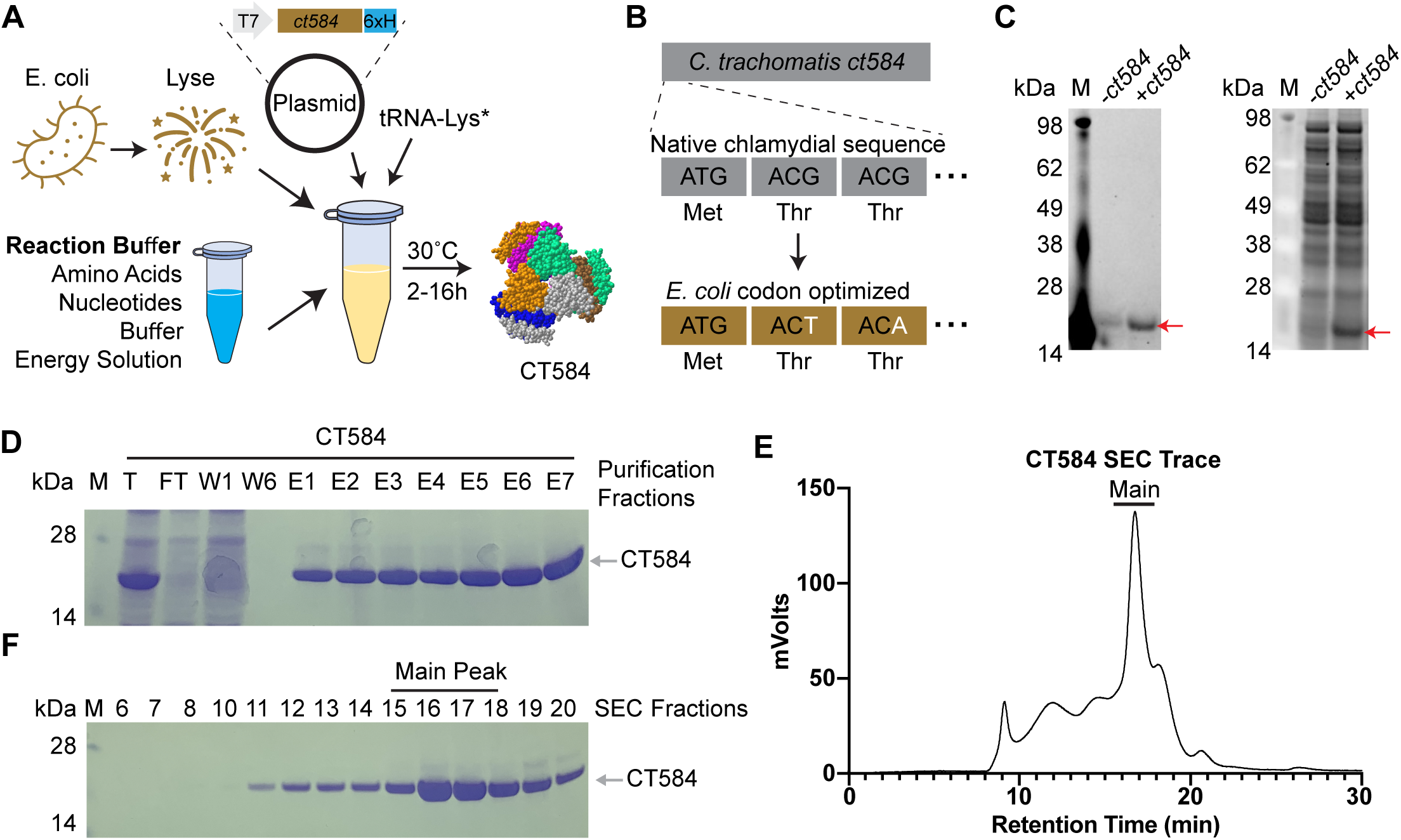
CT584 can be produced and purified using cell-free protein synthesis. **A)** Schematic of the cell-free protein synthesis (CFPS) approach to protein production. **B)** Schematic of DNA codon optimization performed on chlamydial *ct584*. **C)** SDS-PAGE gel depicting fluorescent image of bodipy labeled tRNA-Lys incorporation into CT584 (left) and sypro ruby total protein stain (right). **D)** Coomassie blue stained SDS-PAGE gel showing affinity typical affinity purification fractions for CT584. M: Marker, T: Total, FT: Flow-through, W: Wash, E: Elutions **E)** Smoothed histogram of size exclusion chromatography (SEC) retention times for affinity purified CT584. **F)** Coomassie blue stained SDS-PAGE gel of SEC elution fractions. Fractions pooled and used for further experiments are indicated (“Main Peak”).

To remove impurities from the CFPS lysates, we purified CT584 using a combination of affinity chromatography and size exclusion chromatography (SEC). His-tag and Ni^+2^-NTA affinity chromatography enriched CT584 away from other cell-free lysate proteins (Figure 1D), however commonly seen His-tag purification contaminants from *E. coli* lysates^30^ could potentially pose issues for further immunization studies and confound results. To further enrich and purify CT584, we concentrated it and performed SEC on the sample, leading to >99% pure protein, giving a final yield of >1.5mg from a 1mL scale CFPS reaction (Figure 1E-1F).

### High-speed atomic force microscopy reveals CFPS produced CT584 has a native-like conformation

CT584 has been shown to exist in a hexameric structure^31^, and since CFPS reactions typically do not include many of the native chaperones and other protein folding machinery present in the original host cell, whether CFPS-made CT584 retains that structure was an open question. Our attempts to generate full-length CT584 protein crystals for structural studies were unsuccessful. However, to confirm our cell-free produced CT584 resembled the crystallized truncated CT584 structure, we performed high-speed atomic force microscopy (HS-AFM) to visualize its geometry and gross structure. Purified CT584 was adsorbed onto a mica surface and imaged in buffer solution with an AFM tip with a nominal probe size of 5nm. We were able to observe individual protein units bound to the mica surface, which formed a globular shape with some sub-structure consistently visible in the AFM images. When we used the AFM images to estimate the protein volume, we obtained a value of 563.6±123.6nm^3^ (Figure 2A).

**Figure 2.**
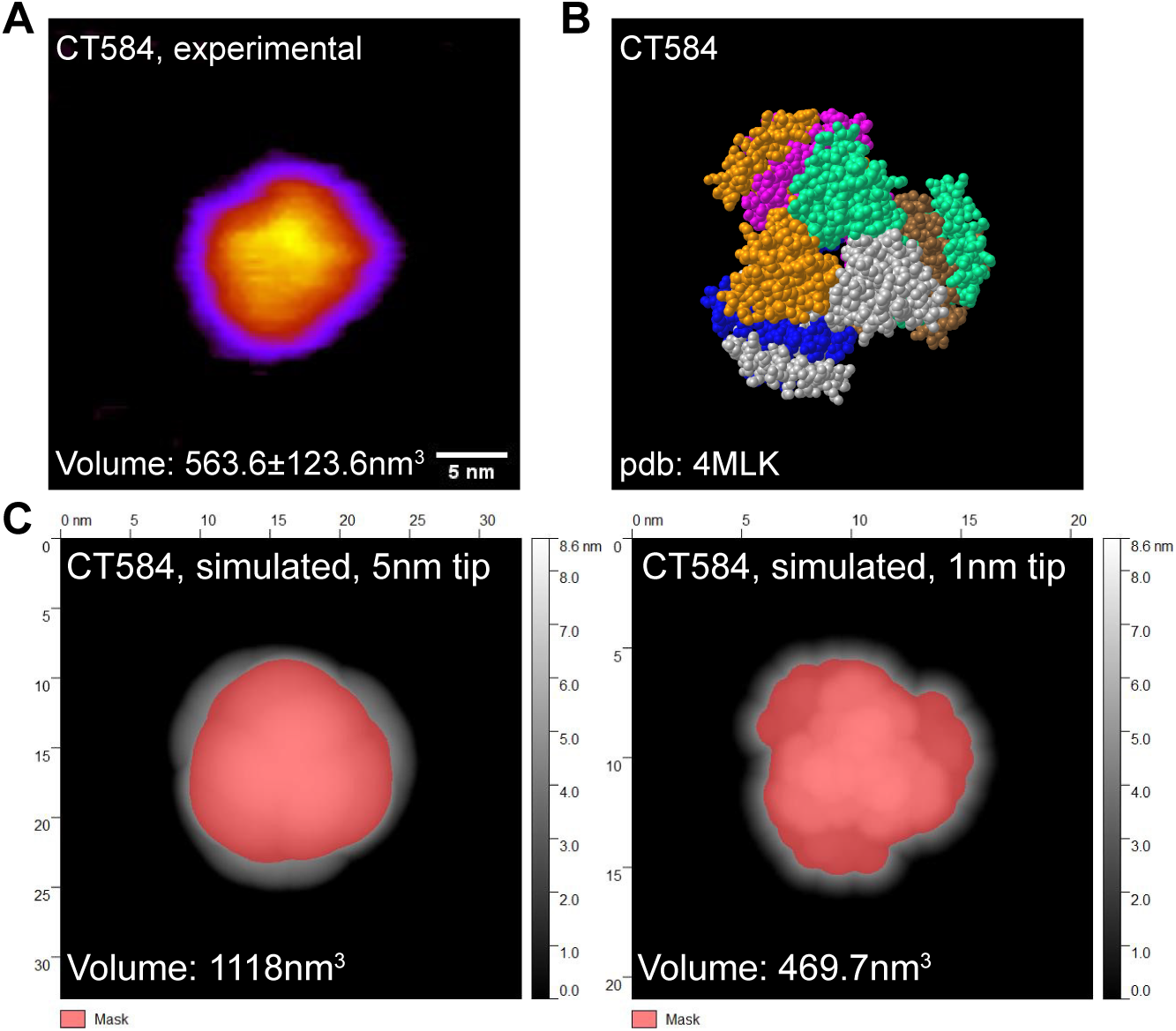
CFPS produced CT584 retains a native-like conformation. **A)** Atomic force microscopy (AFM) image of CT584. Scale bar: 5nm. **B)** Space-filling model of published CT584 crystal structure (PDB: 4MLK). **C)** Simulated AFM images using the CT584 crystal structure and a 5nm (left) and 1nm (right) probe size. Highlighted pink areas show coverage.

From the known crystal structure of truncated CT584^31^ (PDB: 4MLK, Figure 2B), we calculated the hypothetical volume of a CT584 hexamer using different AFM tip sizes as a model parameter (Figure 2C-2D). Using a nominal AFM tip size of 5 nm produced a poor fit (volume estimate of 1118nm^3^) for several reasons. First, the simulated AFM image assumed a perfectly rigid protein which does not deform like a physical sample would. Second, because of the small physical dimensions of the protein, it is likely imaged with individual asperities on the tip surface that can be significantly sharper than the nominal AFM tip size leading to overestimates of volume. Increasing the resolution by simulating a 1nm tip size produced a good fit with the volume estimated at 469.7nm^3^, which was close to our experimentally measured protein volume, suggesting that cell-free produced full-length CT584 may indeed exist in a similar multimeric conformation to the truncated CT584 crystal structure.

### CT584 is immunogenic in mouse models and generates an IgG antibody response

As a putative needle-tip protein of the T3SS, we hypothesized that CT584 could act as a suitable antigen for immunization against chlamydial infection. Both mucosal and systemic routes of immunization have been reported for recombinant protein antigens against chlamydia^6^, and *a priori*, it is difficult to know which route will perform better. To investigate the utility of CT584 as a vaccine antigen and determine the effect of delivery route, we designed two separate but related experiments. We injected 20µg of CT584 or PBS adjuvanted with CpG-1826 and Montanide ISA 720 in a 28-day interval prime (i.n. or i.m.) and i.m. boost, followed by an i.n. challenge regime using female BALB/c mice (Figure 3A). Due to the incompatibility of Montanide for intranasal application, the i.n. prime was only adjuvanted with CpG-1826. Recombinant MOMP (rMOMP) and EBs were also included as positive controls. CpG-1826 has been reported to promote a Th1 response while Montanide ISA 720 induces a mixed Th1/Th2 response^32,33^. Importantly, the combination has been shown to be effective as adjuvants for chlamydial vaccines^34^.

**Figure 3.**
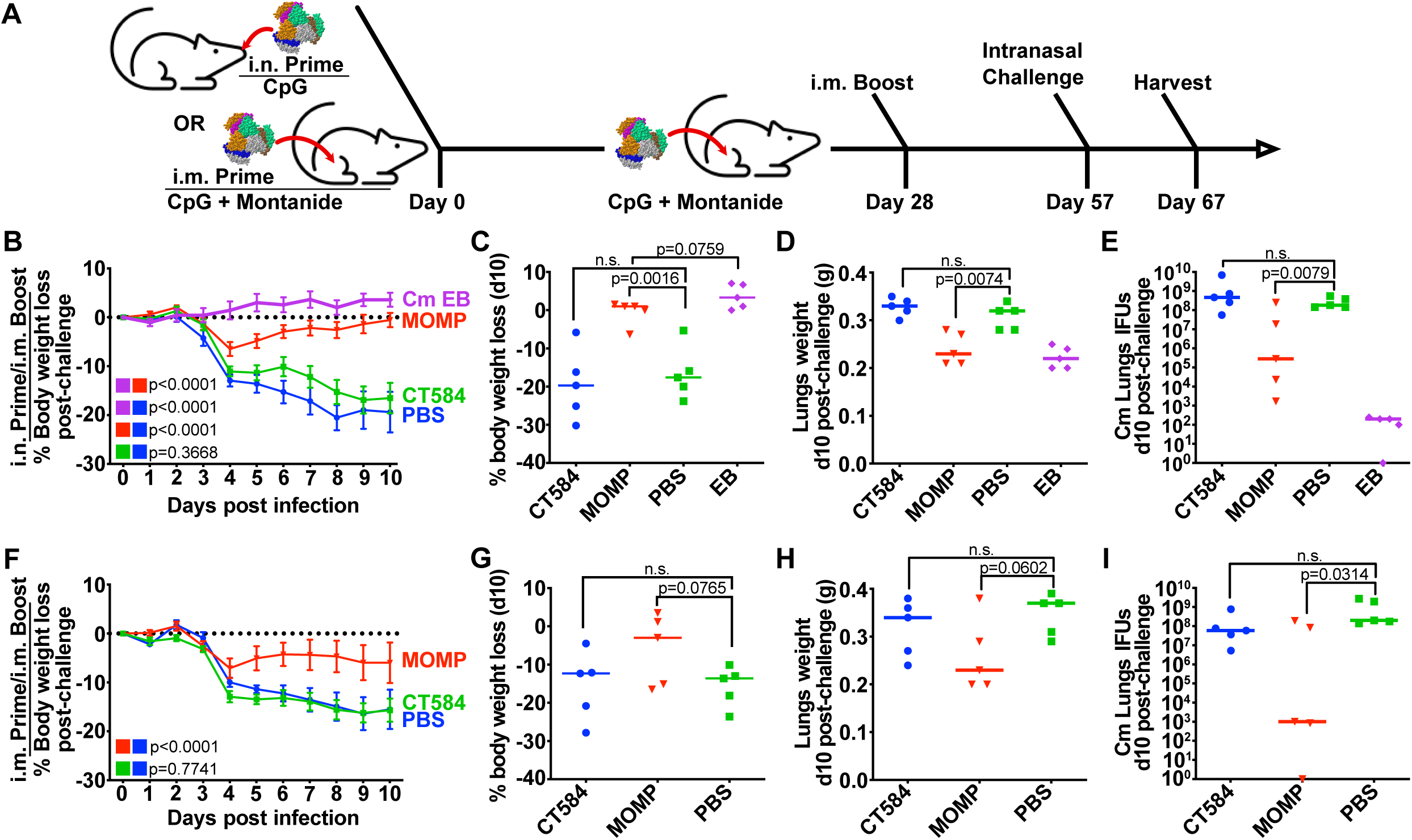
CT584 immunization fails to generate a protective response against an i.n. chlamydial challenge in mice. **A)** Schematic of immunization routes and schedule for challenge studies. **B)** Line graph depicting mean mouse weights +/- s.e.m. after intranasal challenge in i.n./i.m. immunized mice. P-values calculated using two-way repeated measures ANOVA. **C)** Scatter plot showing percent body weight loss at day 10 post-challenge in i.n./i.m. immunized mice. Line indicates median. P-values calculated using Student’s t-test. **D)** Scatter plot showing lung weights at day 10 post-challenge in i.n./i.m. immunized mice. Line indicates median. P-values calculated using Student’s t-test. **E)** Scatter plot showing number of Cm IFUs in mouse lungs at day 10 post-challenge in i.n./i.m. immunized mice. Line indicates median. P-values calculated using Mann-Whitney U test. **F)** Line graph depicting mean mouse weights +/- s.e.m. after intranasal challenge in i.m./i.m. immunized mice. P-values calculated using two-way repeated measures ANOVA. **G)** Scatter plot showing percent body weight loss at day 10 post-challenge in i.m./i.m. immunized mice. Line indicates median. P-values calculated using Student’s t-test. **H)** Scatter plot showing lung weights at day 10 post-challenge in i.m./i.m. immunized mice. Line indicates median. P-values calculated using Student’s t-test. **I)** Scatter plot showing number of Cm IFUs in mouse lungs at day 10 post-challenge in i.m./i.m. immunized mice. Line indicates median. P-values calculated using Mann-Whitney U test.

Serum and mucosal anti-EB titers were determined by ELISA the day before the *Cm* challenge (Table 1). In both i.n. and i.m. regimes, CT584 immunized mice generated an IgG response greater than PBS immunized groups, although the response was weaker compared to rMOMP or EB immunized groups. Antibody titers from CT584 i.n./i.m. immunized mice showed a skew towards IgG2a vs IgG1, indicating a more Th1-biased immune response, while CT584 i.m./i.m. immunized mice generated a more balanced Th1/Th2 immune response. Mucosal IgG and IgA antibody titers from vaginal washes were low or negligible.

**Table 1.** Anti-*Cm* EB titers from immunized mice. Pooled antibody titers from i.n./i.m. (top) or geometric mean titers from i.m./i.m. immunized mouse sera (bottom) are shown along with vaginal wash titers.

| i.n./i.m. Immunization group | Pooled Anti-EB serum dilution titer |  |  | Pooled Vaginal Wash Anti-EB |  |
| --- | --- | --- | --- | --- | --- |
|  | IgG1 | IgG2a | IgG2a/IgG<sub>1</sub> | IgG | IgA |
| CT584 | 800 | 3,200 | 4 | 160 | <10 |
| Cm rMOMP | 51,200 | 51,200 | 1 | 640 | 80 |
| Live Cm EB (i.n) | 1,600 | 25,600 | 16 | 640 | 1,280 |
| PBS | <100 | <100 | - | <10 | <10 |
| i.m./i.m. Immunization group | Anti-EB serum dilution titer |  |  | Vaginal Wash Anti-EB titer |  |
|  | IgG1 | IgG2a | IgG2a/IgG<sub>1</sub> | IgG | IgA |
| CT584 | 3,676 (1,600-6,400) | 3,200 (200-25,600) | 0.9 | 80 | <10 |
| Cm rMOMP | 38,802 (6,400-204,800) | 178,289 (102,400-409,600) | 4.6 | 1,280 | 20 |
| PBS | <100 | <100 | - | <10 | <10 |

### CT584 immunization is not protective against an intranasal *C. muridarum* challenge

To test whether the immune response generated with CT584 is protective, immunized mice were challenged intranasally with 1x10^4^ IFU of *Cm* four weeks after boost (Figure 3A). Mice body weights were tracked for ten days post-challenge before euthanasia and examination of lungs’ weights and determination of the number of *Cm* IFUs.

Both EB and rMOMP immunized groups exhibited significant differences to the PBS control group, showing lower body weight loss (Figure 3B-C, 3F-G), lower lungs’ weights (Figure 3D, 3H), and lower IFUs recovered from the lungs (Figure 3E, 3I). Although the rMOMP immunized groups showed an initial rapid weight loss from days 2-4 after challenge, they recovered to initial body weights by day 10. In contrast, the CT584 immunized mice performed similarly to the PBS negative control group, showing rapid body weight loss, increased lungs’ weights, and higher lungs’ IFUs recovered compared to the EB and rMOMP positive control groups.

To determine whether the CT584-induced antibodies were antigen-specific, we used pooled immune sera from mice as the primary antibody in a western blot of purified CT584 protein (Figure 4A). As expected, the CT584 immune sera detected a single robust band at the correct molecular weight while the PBS immune sera did not detect any protein. Unexpectedly, sera from mice immunized with *Cm* EBs also failed to detect CT584. Running the converse experiment, we blotted whole *Cm* EB lysates onto membranes and probed with the same pooled immune sera (Figure 4B). Although CT584 immune sera could detect CT584 in EB lysates, EB immune sera could not, raising the question of whether CT584 antigens are accessible or immunogenic in EBs for antibody generation or are present in too low quantities.

**Figure 4.**
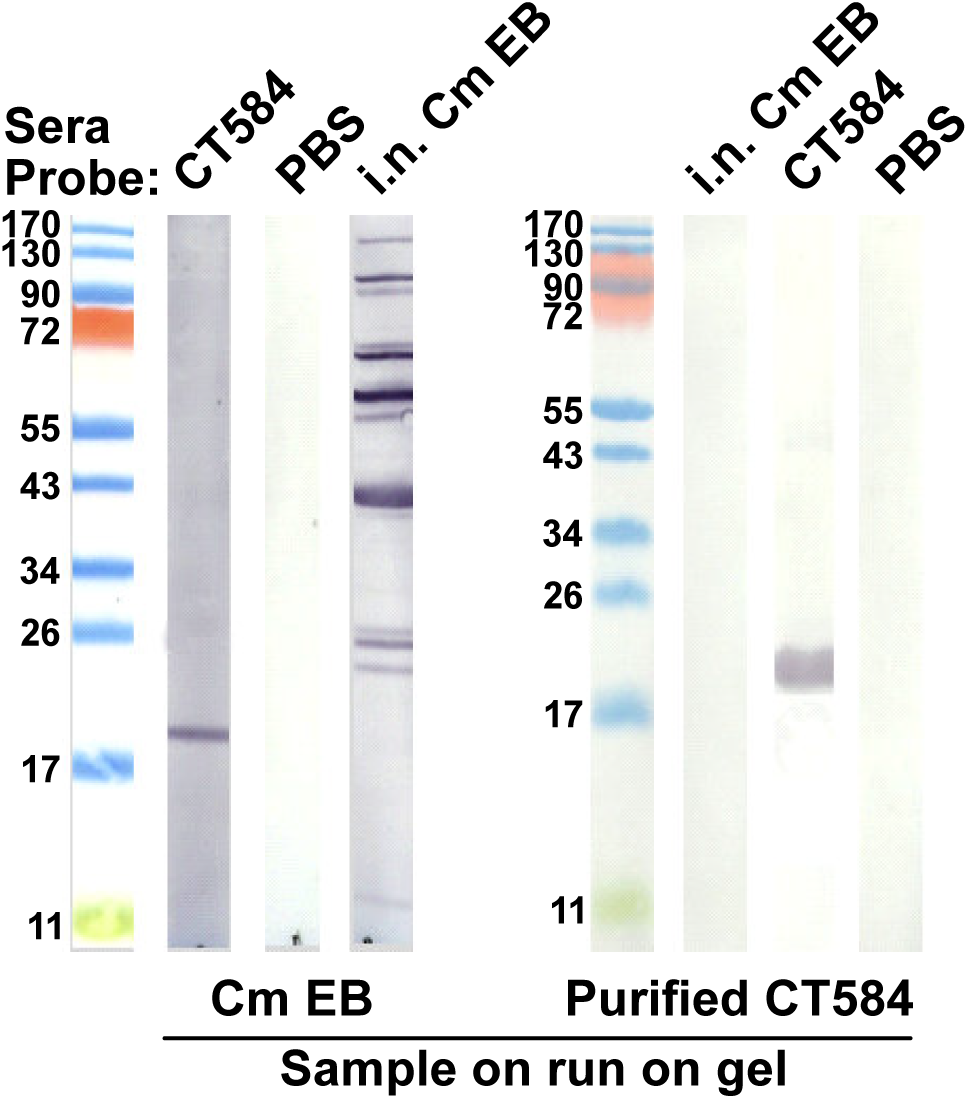
CT584 immune sera detects the recombinant and native protein but *Cm* EB immune sera does not. **A)** Western blots of purified recombinant CT584 probed with the serum of mice immunized with: Lane 1 – recombinant CT584; Lane 2 – PBS; Lane 3 – i.n. *Cm* EB. **B)** Western blots of *Cm* EB lysates probed with the serum of mice immunized with: Lane 1 - i.n. live *Cm* EB; Lane 2 - recombinant CT584; Lane 3 – PBS.

Since adjuvant choice in formulations can have a significant influence on vaccine effectiveness and outcomes, we also investigated the immune response generated towards CT584 using CAF01 and R848 adjuvants in a 28-day interval i.m. prime-boost-boost regimen in BALB/C mice (Figure 5A). These adjuvants have been described to stimulate Th1/Th2 balanced and Th2-biased, respectively, immune responses^35,36^. Consistent with our previous results, we observed an IgG response induced by CT584 immunization. Furthermore, we observed elevated IgG1 vs. IgG2a antibody levels, suggesting a Th2 favored response (Figure 5B-C). To determine what T-cell immune responses were induced, we examined the CD4 and CD8 T cell population (Figure 5D-E) extracted from the draining lymph nodes and spleens of immunized mice after euthanasia and performed *ex vivo* antigen restimulation assays. We assessed markers such as IFN-γ (Figure 5F-G, left) and TNF-α (Figure 5F-G, right), however, there were no significant differences between CT584 immunized mice and PBS controls for any marker.

**Figure 5.**
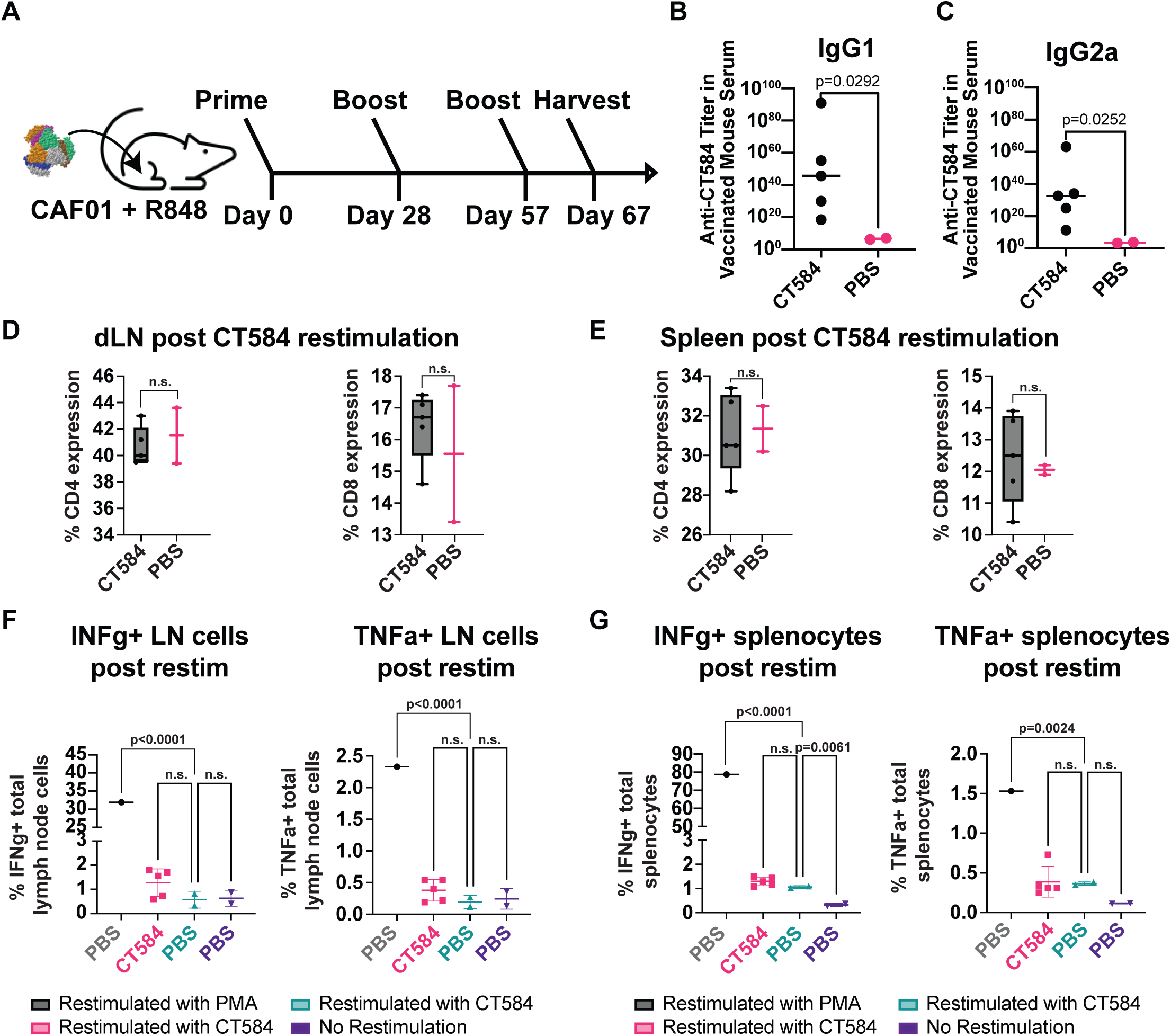
CT584 + CAF01 and R848 induces antigen-specific antibodies but not T cell-based responses. **A)** Schematic of i.m./i.m. immunization routes and schedule for CT584 with CAF01/R848. **B)** Scatter plot showing CT584 specific IgG1 antibodies in mouse immune sera. Line indicates median. P-values calculated using Welch’s t-test. **C)** Scatter plot showing CT584 specific IgG2a antibodies in mouse immune sera. Line indicates median. P-values calculated using Welch’s t-test. **D)** Box and whiskers plot showing CD4 (left) and CD8 (right) expression in T cell populations from the draining lymph nodes of immunized mice. P-values calculated by Student’s t-test. **E)** Box and whiskers plot showing CD4 (left) and CD8 (right) expression in T cell populations from the spleen of immunized mice. P-values calculated by Student’s t-test. **F)** Scatter plot showing percentage of lymph node immune cells expressing IFN-γ (left) and TNF-α (right) after ex vivo restimulation with antigen. Line indicates median. P-values calculated by one-way ANOVA. **G)** Scatter plot showing percentage of splenocytes expressing IFN-γ (left) and TNF-α (right) after ex vivo restimulation with antigen. Line indicates median. P-values calculated by one-way ANOVA.

## Discussion

There is a growing need for access to vaccines in the world, especially against *Ct*, where currently antibiotics are the only treatment modality. Although it is the most prevalent bacterial STI, its health consequences disproportionately impact poor and rural regions worldwide, where myriad logistical challenges towards detection and administration of aid remain^37^. Synthetic biology techniques such as CFPS can potentially address these gaps, promising the ability to rapidly generate protective antigens at the point of use. CFPS lysates are amenable towards lyophilization and reconstitution in a just-add-water system, making them ideal for low-cost low requirement solutions^38,39^. Demonstrably, pilot studies generating an on-demand vaccine against *Francisella tularensis* have been performed^40^. Due to the inherent flexibility of CFPS systems, it is also possible to produce and solubilize even challenging chlamydial antigens, such as the integral membrane protein MOMP, for immunization studies where it was able to generate a protective response in mice^41,42^. Taking advantage of this flexibility, we used CFPS to produce and characterize the effects of a T3SS protein on protection against *Cm* infection.

T3SS function is highly conserved across chlamydial species and is required for infection^43^, making it an attractive target for potential vaccines against *Ct*. While chlamydial gene expression can vary over the course of infection due to its distinctive bi-phasic life cycle, the T3SS is present in both EBs and RBs^11^, suggesting a potential vaccine against T3SS components could be useful at all stages of infection. T3SS needle tip antigens from other organisms such as *Y. pestis*^44,45^ and *Salmonella enterica*^46,47^ have been successfully used as vaccine antigens to protect against lethal challenge in mice and non-human primate models. Because CT584 was identified as a putative needle-tip protein in the chlamydial T3SS, we reasoned that it could be an attractive vaccine antigen candidate.

We produced and purified recombinant CT584 using CFPS and evaluated it for usage as a protective chlamydial vaccine antigen. We confirmed that CFPS-produced CT584 resembles its known structure (PDB: 4MLK^31^) using AFM data and simulations. However, we did not see any protection from mice immunized with 20µg of CT584, neither through i.m. nor i.n. routes, after i.n. challenge with *Cm* EBs. The CT584 immunized groups showed no difference to the PBS control, resulting in significant weight loss, increased lung weights indicative of ongoing inflammation, and increased lungs’ IFUs.

CT584 has been incorporated into two potential chlamydial vaccine formulations to date. As a recombinant protein fusion with other T3SS proteins, CopB and CopD, CT584 elicited a reduction in bacterial shedding in both *Cm* and *Ct* genital tract challenge models^15,16^. However, when using peptides designed from B-cell specific epitopes conjugated to a bacteriophage Qβ-based virus-like-particle (VLP) platform^48^, results were more mixed. Although initial experiments showed a reduction in *Ct* burden upon transcervical challenge, repeated experiments did not reach statistical significance. It is important to note that in the current study as well as the three studies described above, vaccination elicited robust antigen-specific antibody titers. While it is widely accepted that an effective chlamydia vaccine requires the induction of a cell-mediated immune response, the contribution of humoral immunity is less clear^49,50^. However, pre-incubation of chlamydial EBs with immune sera from VLP-CT584 immunized mice from Webster et al.^48^ before transplantation into naïve mice resulted in a significant reduction in upper genital tract bacterial burden compared to sera from VLP alone, highlighting the potential role antibodies may play in protection against chlamydial infection.

This begs the question: Although pre-opsonized EBs showed a marked reduction in bacterial load, and presumably, infection efficiency, why are the antibodies generated against CT584 generally insufficient for protection against challenge with live EBs? Speculatively, it is possible that these neutralizing antibodies are not present in sufficient titers on the mucosal surfaces – indeed we observe significantly less IgG present in the vaginal washes compared to sera, as well as negligible IgA production, in our experiments. Alternatively, there may be issues with CT584 antigen presentation *in vivo*. While CT584 immune sera could detect CT584 protein in EB lysates or purified samples through WB, the converse was not true. EB immune sera could not detect CT584 in any condition, suggesting that CT584 antigen in EBs may be masked, not present on the cell surface, or otherwise not available for antibody generation. Irrespective of the cause, our data suggests that CT584 alone may not be suitable as a vaccine candidate.

Although CT584 by itself did not elicit a protective immune response against *Cm* respiratory challenge in our formulations, the T3SS in general may still be a promising target. T3SS function is crucial for chlamydial infection, and immunization using fusions of the terminal translocon pore proteins CopB and CopD showed encouraging results in both *Cm* and *Ct* mouse models^15,16^. Because fusion proteins, especially those of integral membrane proteins, may not be in the properly folded state, it may be possible to improve upon that antigen. Combining CFPS with nanodisc^41^ technology would allow for the de novo synthesis of intact CopB and CopD pore complexes in membrane discs, mimicking their native structure, and could make for an ideal T3SS antigen for future studies.

## Materials and Methods

### Plasmids and sequences

The amino acid sequence from *Ct* CT584 (PDB: 4MLK_A) was reverse-translated, codon optimized for expression in *E. coli*, and synthesized by Genscript with a short linker (AAALE) sequence before a C-terminal 6x histidine affinity purification tag. The gene fragment was cloned into the NdeI-BamHI sites in pIVEX2.3d for T7 polymerase-based expression. Amino acid sequences from *C. muridarum (Cm)* TC0873 (Uniprot Q9PJF6) and C. trachomatis (*Ct)* CT584 (Uniprot O84588) were downloaded from Uniprot using accession ID’s Q9PJF6 and O84588, respectively. *Cm tc0873* and *Ct ct584* gene sequences were downloaded from NCBI through ChlamBase^51^ using accession ID’s NC_002620.2 (locus: 1010962..1011513, + strand) and NC_000117.1 (locus 657866..658417, + strand), respectively. Sequence homology was compared using the Smith-Waterman algorithm in SnapGene, software version 7.2.

### Stocks of C. muridarum

*C. muridarum* (strain Nigg II; American Type Culture Collection) was grown in HeLa-229 cells using high glucose Dulbecco’s medium, plus cycloheximide (1 µg/ml) and gentamycin 10 µg/ml, without fetal bovine serum. Elementary bodies (EB) were purified and stored in sugar phosphate glutamate buffer (SPG) at -80°C as described^8^. The number of *C. muridarum* inclusion forming units (IFU) in the stock was assessed in HeLa-229 cells using immune-peroxidase staining with a *Cm*-MOMP specific mAb (MoPn-40) produced in our laboratory^52^.

### Cell-free protein synthesis of CT584 and purification strategies

CT584 protein was produced using cell-free protein synthesis kits in a continuous-exchange cell-free (CECF) device (Biotech Rabbit, #BR1400201) according to manufacturer protocols at 1mL scale. Briefly, lyophilized E. coli lysate, reaction buffer, amino acids, and feed buffer were reconstituted using the supplied reconstitution buffer supplemented with an EDTA-free protease inhibitor tablet (Roche, #5892791001). The individual components were combined, and CT584-His in pIVEX2.3d plasmid was added to the reaction mixture at 15µg/mL before transferring into the CECF device. BODIPY-FL labeled tRNA-Lys (Promega L5001) were included in initial test reactions at 1:200 v/v scale. The reaction was incubated at 30°C for 16 hours while shaking at 300rpm in a bacterial floor shaker.

After incubation, the CFPS reaction mixture was removed from the CECF device and purified using Ni^+2^-NTA affinity chromatography. 500uL of Ni+2-NTA beads (Roche, #5893682001) were packed onto a 12mL column and equilibrated into six packed bead volumes of equilibration buffer (50mM Tris, 300mM NaCl, and 10mM imidazole, pH 8.0). The CFPS reaction mixture was applied to the column and allowed to incubate at 4°C for 1 hour while mixing on a nutator. Afterwards, the column was washed with twelve packed bead volumes of wash buffer (50mM Tris, 300mM NaCl, and 20mM imidazole, pH 8.0). CT584 was eluted, first with 1.8mL of elution buffer (50mM Tris, 300mM NaCl, and 250mM imidazole, pH 8.0), and then 300uL of elution buffer containing 500mM imidazole.

Elution fractions 2-7 were pooled and concentrated to 200uL using a 10k MWCO filtration column (Amicon, #UFC501096). The concentrated protein was injected into a Superdex 200 Increase 10/300 GL (GE Healthcare 28-9909-44) size exclusion column chromatography system (Waters). Peak fractions were collected and pooled for further downstream analysis.

### HS-AFM data collection

HS-AFM images of Dyn-lipid nanotube complexes were acquired in tapping mode at room temperature using a HS-AFM instrument (RIBM, Japan) equipped with ultra-short AFM cantilevers Olympus, BL-AC10FS) with EBD tip (radius < 5 nm). This instrument utilizes dynamic proportional-integral-differential (PID) controller^53,54^ to eliminate the probe “parachuting” artifacts from images and reduce the tip-sample forces. HS-AFM fluid cell was filled with 120 μL buffer solution (20 mM HEPES (pH 7.2), 150 mM KCl, 1 mM DTT, 1mM EDTA, 1 mM EGTA) and the constant volume was maintained with a custom-built perfusion system. In a typical experiment we collected 128 pixel × 128 pixel images from a 50 nm × 50 nm area at a scan rate of 3 frames per second.

### HS-AFM data processing and analysis

Raw HS-AFM image data was converted to ImageJ (developed by Wayne Rasband, National Institutes of Health, Bethesda, MD) stacks using custom software^55^. The line plots were generated by a custom IgorPro 8 software (WaveMetrics, Lake Oswego, OR, USA). The simulated AFM images were generated by using open-source code^56^. The volume of protein oligomer from experimental data and simulated AFM images were calculated by using Gwyddion software^57^.

### Vaccination of female BALB/c mice

For challenge studies, four-to-five-week-old female BALB/c (H-2d) mice (Charles River Laboratories) were obtained and housed in the University of California, Irvine, vivarium and experiments carried out according to the approved Institutional Animal Care and Use Committee (IACUC) protocol. Mice were vaccinated with CT584 (20μg/mouse/immunization) using two different regimens: 1) Intranasal (i.n.) prime and intramuscular (i.m.) boost in the quadriceps muscle at a four-week interval; and 2) i.m. prime and boost four weeks apart in the quadriceps muscle. The following adjuvant combinations were used: CpG-1826 (Tri-Link) (10 µg/mouse/immunization) + Montanide ISA 720 VG (Seppic Inc.)^52^ at 70% (v/v) of the final dose. For i.n. immunization, only CpG-1826 (10 µg/mouse/immunization) was used. For all i.m. vaccinations, Montanide ISA 720 VG was mixed with CT584 and CpG-1826 using a vortex (Fisher Scientific). The formulation was vortexed for one minute followed by one minute rest on ice. The cycle was repeated five times. A negative immunization control received PBS.

For alternative adjuvant studies, 5-week-old female BALB/c (H-2d) mice were obtained and housed in the Lawrence Livermore National Laboratory vivarium and experiments carried out according to the approved IACUC protocol. Mice were vaccinated i.m. with CT584 (10µg/mouse/immunization) adjuvanted with CAF01 (50µg) and R848 (Invivogen) (20µg) in a four-week interval prime-boost-boost regime. CAF01 is a liposomal adjuvant^58^ prepared by mixing dimethyldioctadecylammonium (DDA) bromide (Avanti) and a,a’-trehalose 6,6’-dibehenate (TDB) (Avanti). Briefly, DDA is mixed with TDB in a 5:1µg ratio, dried under a nitrogen stream with continuous agitation, and desiccated in a vacuum overnight before resuspension in water. After resuspension, the liposomes are heated at 57°C for 20 minutes while shaking and then sonicated for 10 minutes before addition to the antigen.

### Intranasal challenge with *Cm* EBs and evaluation of the course of the infection in mice

Anesthetized mice were challenged i.n. with 10^4^ IFU of *Cm* four weeks after the second immunization^59^. Daily body weight changes were assessed for 10 days post-challenge (d.p.c.) when mice were euthanized, their lungs weighed and homogenized (Seward Stomacher 80; Lab System) in 5 ml of SPG. To establish the number of *Cm* IFU, six serial dilutions of the lungs’ homogenates were used to infect Hela-229 cells grown in 48 well plates. Following incubation for 30 h at 37°C in a CO2 incubator, the IFU were visualized with mAb MoPn-40, and counted using a light microscope^60^. The limit of detection (LD) was < 50 *Cm* IFU/lungs/mouse.

### Serum Antibody Titers

CT584-specific antibody titers were quantified by ELISA as previously described^42^. Briefly, sera from PBS or sham immunized mice were used as negative controls. 96-well, high binding plates (Corning™ 3690) were coated with 100 µL of CT584 at a concentration of 10 µg of protein/mL and incubated overnight at the 4°C. Serum was added to each well in 2-fold serial dilutions after washing and blocking for nonspecific binding. After overnight incubation, the plates were washed and incubated with one of the following secondary antibodies: Horseradish peroxidase-conjugated (HRP) goat anti-mouse IgG antibodies (KPL 5220-0460), HRP rat anti-mouse IgG1 (BD Biosciences 559626), and HRP rat anti-mouse IgG2a conjugate (BD Biosciences 553391). The binding was measured using an BioTek (Agilent) plate reader or Labsystems Multiscan RC (ThermoFisher) at an optical density (OD) of 405 nm. The geometric mean titers are expressed as the reciprocal of the dilution.

### Tissue harvest and single cell isolation

Mouse sera was collected 3 weeks after the last boost by cardiac puncture. Spleens and popliteal draining lymph nodes were also harvested. Single cells from spleens and draining lymph nodes were isolated by gently crushing the tissues individually through a sterile 70 µm cell strainer while passing sterile HBSS (Gibco™ 14025092) supplemented with 2% heat inactivated FBS (Gibco™ A5669801) to wash down the cells. Following centrifugation at 1500 RPM for 5 minutes at 4°C, red blood cells from the splenocytes and draining lymph node cells were lysed by resuspending the cell pellets with 3 mL of ammonium-chloride-potassium (ACK) lysing buffer (Gibco™ A1049201) and incubation for 5 minutes at room temperature, followed by dilution of the mixture with 27 mL of the wash buffer (HBSS + 2% FBS) to stop the reaction. The lysed cells were pelleted and resuspended with culture media (RPMI 1640 medium (Gibco™ 11875093) supplemented with 10% FBS) for *ex vivo* restimulation.

### SDS-PAGE/WBs

Aliquots of total cell-free reaction or purified CT584 SEC fractions were diluted into 4x NuPAGE LDS Sample Buffer (ThermoFisher NP0007) and 10x NuPAGE Sample Reducing Agent (ThermoFisher NP0009) before heat denaturing. Samples were loaded onto 4-12% NuPAGE Bis-Tris gradient gels (ThermoFisher NP0323BOX) and run at 200V for 35 minutes. Gels were stained with SYPRO-Ruby Protein Gel Stain (ThermoFisher S12000) according to manufacturer directions and imaged on an Odyssey Fc Imager (Li-Cor).

Western blot was performed to detect CT584-specific antibody binding. Briefly, either the recombinant CT584 or purified *Cm* EB’S were solubilized in SDS sample buffer (Novex, Life Technologies) in the presence of reducing agent (10mM DTT), heated in a water bath at 95-100°C for 5 min and run on 10% SDS-PAGE gels. After electrophoresis, the resolved protein bands were transferred onto nitrocellulose membranes and the membrane blots were detected with mouse sera collected a day before challenge as the primary antibody. The resulting antibody binding was probed with HRP goat anti-mouse immunoglobulins (IgG, IgA, IgM) (Cappel, MP Biomedicals) as the secondary antibody and visualized with chloronaphtol substrate.

### *Ex vivo* restimulation

Splenocytes and lymph node cells were seeded in 24 well plates at 1 million cells/well/sample and cultured with CT584 at a concentration of 10 μg of protein/mL in culture media supplemented with 1x monensin, a protein that enhances intracellular cytokine staining signals by blocking transport processes during cell activation.

### Flow Cytometric Analysis

A 10-color T cell panel was used for flow cytometry to delineate IFNγ and TNFα expressing CD4+ and CD8+ T cells, and to evaluate changes in both CD4 and CD8 T cell subsets. Briefly, cultured splenocytes and lymph node cells were washed and resuspended in FACS staining buffer (BioLegend 420201) containing 1 μg of purified Rat Anti-Mouse CD16/CD32 (Mouse BD Fc Block) (BD Pharmingen 553141) and were kept on ice throughout staining. Cell surface staining antibodies (PI, CD45, CD3, CD4, CD8, CD62L, and CD44) were added and cells were incubated on ice for 45 min. Cells were washed and then fixed and permeabilized using the True-Nuclear™ Transcription Factor Buffer Set (BioLegend 424401). Intracellular staining antibodies (IFNγ, TNFα, and FoxP3) were added as recommended in the permeabilization buffer. Following incubation, cells were washed and resuspended in FACS staining buffer for analysis on a BD FACSCelesta Cell Analyzer (BD Biosciences, San Jose, CA). Flow cytometry data was analyzed using FlowJo v10 (BD Biosciences, San Jose, CA). Antibodies against cell-surface and intracellular markers were obtained from either BD Biosciences or BioLegend.

### Statistical analyses

The Student’s t test was employed to evaluate differences between changes in body weight at day 10 post-challenge, lungs’ weights, and CD4 and CD8 T-cell populations. Two-way repeated measures ANOVA with Sidak’s multiple comparison test was employed to compare changes in mean body weight over the 10 days of observation following the *Cm* i.n. challenge. The Mann-Whitney U Test was used to compare the number of *Cm* IFU in the lungs. Welch’s t-test was used to compare antibody titers. ANOVA using Dunnett’s multiple comparisons test was used to compare IFN-γ and TNF-α levels. A P value of < 0.05 was considered to be significant. A P value of < 0.1 was regarded as approaching significance.

## Author Contributions

Conceptualization, S.H.-P., L.M.d.l.M., and M.A.C.; Formal Analysis, S.H.-P., S.P., A.S., A.A.-O., F.B., M.S., and Y.Z.; Investigation, S.H.-P., S.P., A.S., A.A.-O., Y.Z., S.F.G., M.V.M., A.R., and L.M.d.l.M; Writing – Original Draft Preparation, S.H.-P. and M.A.C.; Writing – Review & Editing, S.H.-P., S.P., A.S., A.A.-O., Y.Z., A.N., A.R., L.M.d.l.M., and M.A.C.; Visualization, S.H.-P., S.P., and A.S.; Supervision, A.N., A.R., L.M.d.l.M., and M.A.C.; Funding Acquisition, A.R., L.M.d.l.M., and M.A.C.

## Acknowledgements

We would like to thank P. D’haeseleer and L. Beckett for their helpful discussions and comments. Work at the Lawrence Livermore National Laboratory was performed under the auspices of the U.S. Department of Energy under Contract DE-AC52-07NA27344.

## Institutional Review Board Statement

These studies were approved by the University of California, Irvine, Institutional Animal Care and Use Committee (IACUC), and Lawrence Livermore National Laboratory IACUC.

---

**Figure S1.**
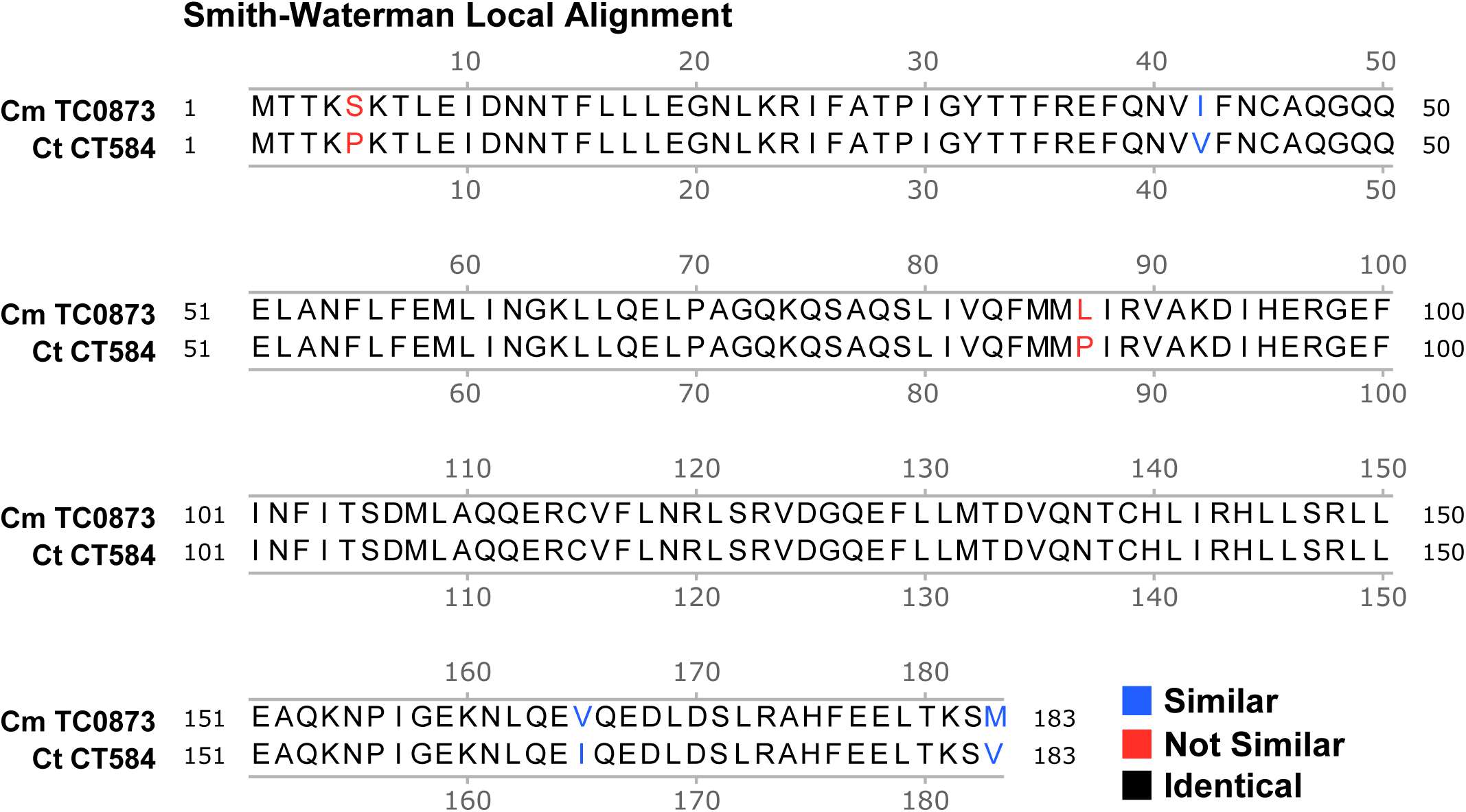
*Cm* TC0873 and *Ct* CT584 are nearly identical. **A)** Smith-Waterman local sequence alignment for *Cm* TC0873 and *Ct* CT584

**Figure S2.**
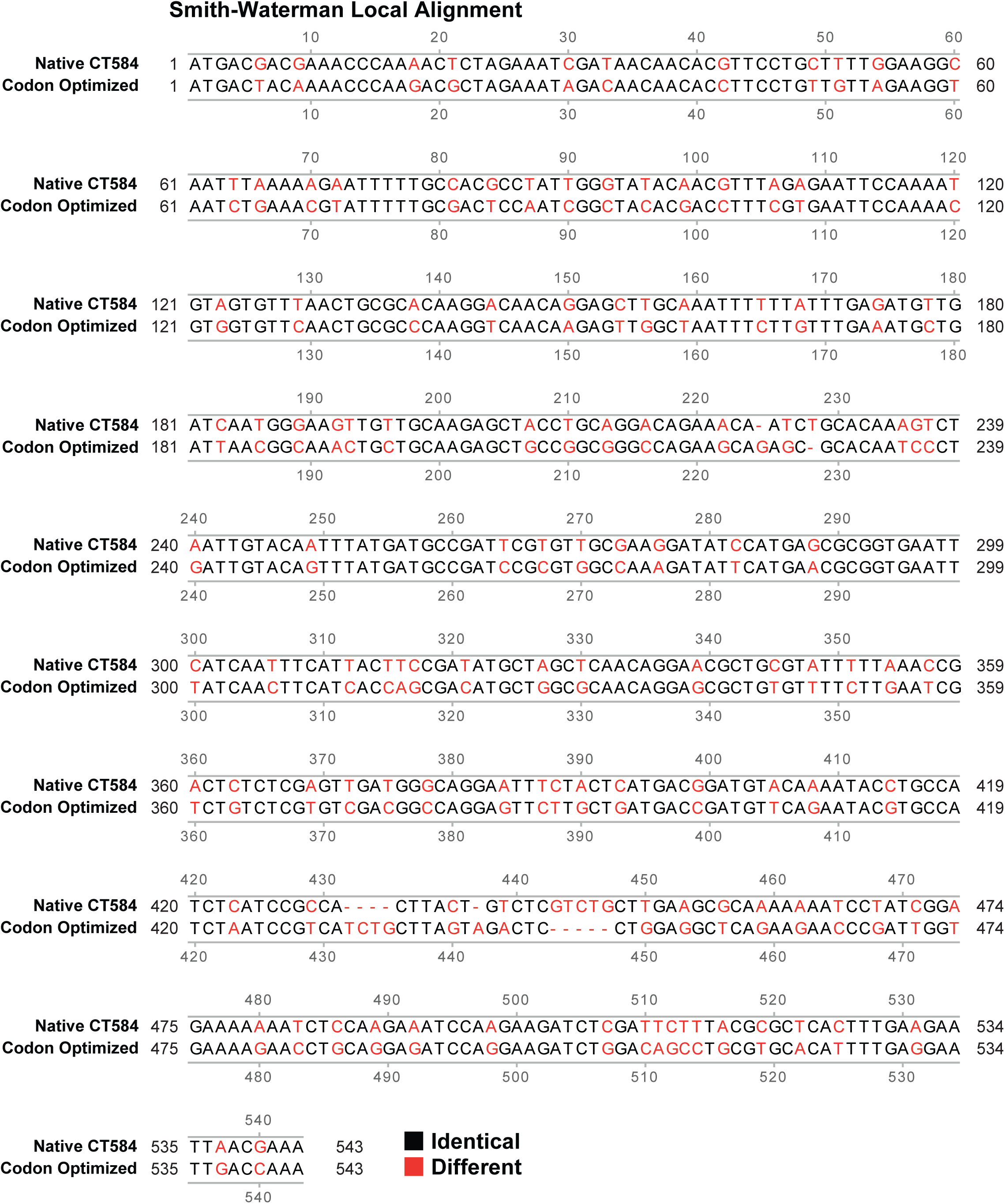
*E. coli* DNA codon optimization for *Ct* CT584. **A)** Smith-Waterman local sequence alignment for the native *Ct* CT584 and *E. coli* optimized DNA sequence

## References

1 Newman, L., et al. Global Estimates of the Prevalence and Incidence of Four Curable Sexually Transmitted Infections in 2012 Based on Systematic Review and Global Reporting. PLoS One 10, e0143304 (2015). 10.1371/journal.pone.0143304

2 den Heijer, C. D. J., et al. Chlamydia trachomatis and the Risk of Pelvic Inflammatory Disease, Ectopic Pregnancy, and Female Infertility: A Retrospective Cohort Study Among Primary Care Patients. Clin Infect Dis 69, 1517–1525 (2019). 10.1093/cid/ciz429

3 Tang, W. et al. Pregnancy and fertility-related adverse outcomes associated with Chlamydia trachomatis infection: a global systematic review and meta-analysis. Sex Transm Infect 96, 322–329 (2020). 10.1136/sextrans-2019-053999

4 Du, M. et al. Increasing incidence rates of sexually transmitted infections from 2010 to 2019: an analysis of temporal trends by geographical regions and age groups from the 2019 Global Burden of Disease Study. BMC Infect Dis 22, 574 (2022). 10.1186/s12879-022-07544-7

5 Pinto, C. N. et al. Impact of the COVID-19 Pandemic on Chlamydia and Gonorrhea Screening in the U.S. Am J Prev Med 61, 386–393 (2021). 10.1016/j.amepre.2021.03.009

6 Peterson, E. M., You, J. Z., Motin, V. & de la Maza, L. M. Intranasal immunization with Chlamydia trachomatis, serovar E, protects from a subsequent vaginal challenge with the homologous serovar. Vaccine 17, 2901–2907 (1999). 10.1016/s0264-410x(99)00131-0

7 Stary, G. et al. VACCINES. A mucosal vaccine against Chlamydia trachomatis generates two waves of protective memory T cells. Science 348, aaa8205 (2015). 10.1126/science.aaa8205

8 Caldwell, H. D., Kromhout, J. & Schachter, J. Purification and partial characterization of the major outer membrane protein of Chlamydia trachomatis. Infection and Immunity 31, 1161–1176 (1981). 10.1128/iai.31.3.1161-1176.1981

9 de la Maza, L. M., Darville, T. L. & Pal, S. Chlamydia trachomatis vaccines for genital infections: where are we and how far is there to go? Expert Rev Vaccines 20, 421–435 (2021). 10.1080/14760584.2021.1899817

10 Nans, A., Kudryashev, M., Saibil, H. R. & Hayward, R. D. Structure of a bacterial type III secretion system in contact with a host membrane in situ. Nat Commun 6, 10114 (2015). 10.1038/ncomms10114

11 Shaw, E. I. et al. Three temporal classes of gene expression during the Chlamydia trachomatis developmental cycle. Mol Microbiol 37, 913–925 (2000). 10.1046/j.1365-2958.2000.02057.x

12 Bailey, L. et al. Small molecule inhibitors of type III secretion in Yersinia block the Chlamydia pneumoniae infection cycle. FEBS Lett 581, 587–595 (2007). 10.1016/j.febslet.2007.01.013

13 Muschiol, S. et al. A small-molecule inhibitor of type III secretion inhibits different stages of the infectious cycle of Chlamydia trachomatis. Proc Natl Acad Sci U S A 103, 14566–14571 (2006). 10.1073/pnas.0606412103

14 Muschiol, S., Normark, S., Henriques-Normark, B. & Subtil, A. Small molecule inhibitors of the Yersinia type III secretion system impair the development of Chlamydia after entry into host cells. BMC Microbiol 9, 75 (2009). 10.1186/1471-2180-9-75

15 Bulir, D. C. et al. Immunization with chlamydial type III secretion antigens reduces vaginal shedding and prevents fallopian tube pathology following live C. muridarum challenge. Vaccine 34, 3979–3985 (2016). 10.1016/j.vaccine.2016.06.046

16 Liang, S. & Mahony, J. B. Intranasal vaccination with a Chimeric Chlamydial Antigen BD584 confers protection against Chlamydia trachomatis genital tract infection. Journal of Vaccines and Immunology 6, 010–017 (2020). 10.17352/jvi.000030

17 Bulir, D. C. et al. Chlamydia Outer Protein (Cop) B from Chlamydia pneumoniae possesses characteristic features of a type III secretion (T3S) translocator protein. BMC Microbiol 15, 163 (2015). 10.1186/s12866-015-0498-1

18 Bulir, D. C. et al. Chlamydia pneumoniae CopD translocator protein plays a critical role in type III secretion (T3S) and infection. PLoS One 9, e99315 (2014). 10.1371/journal.pone.0099315

19 Markham, A. P., Jaafar, Z. A., Kemege, K. E., Middaugh, C. R. & Hefty, P. S. Biophysical characterization of Chlamydia trachomatis CT584 supports its potential role as a type III secretion needle tip protein. Biochemistry 48, 10353–10361 (2009). 10.1021/bi901200y

20 Cowan, C., Philipovskiy, A. V., Wulff-Strobel, C. R., Ye, Z. & Straley, S. C. Anti-LcrV antibody inhibits delivery of Yops by Yersinia pestis KIM5 by directly promoting phagocytosis. Infect Immun 73, 6127–6137 (2005). 10.1128/IAI.73.9.6127-6137.2005

21 Philipovskiy, A. V. et al. Antibody against V antigen prevents Yop-dependent growth of Yersinia pestis. Infect Immun 73, 1532–1542 (2005). 10.1128/IAI.73.3.1532-1542.2005

22 de la Maza, L. M., Pal, S., Khamesipour, A. & Peterson, E. M. Intravaginal Inoculation of Mice with the Chlamydia trachomatis Mouse Pneumonitis Biovar Results in Infertility. Infection and Immunity 62, 2094–2097 (1994). 10.1128/iai.62.5.2094-2097.1994

23 Nigg, C. AN UNIDENTIFIED VIRUS WHICH PRODUCES PNEUMONIA AND SYSTEMIC INFECTION IN MICE Science Jan 995, 49–50 (1942). 10.1126/science.95.2454.49-a

24 Garenne, D. et al. Cell-free gene expression. Nature Reviews Methods Primers 1 (2021). 10.1038/s43586-021-00046-x

25 Batista, A. C., Soudier, P., Kushwaha, M. & Faulon, J. L. Optimising protein synthesis in cell-free systems, a review. Eng Biol 5, 10–19 (2021). 10.1049/enb2.12004

26 Li, X. et al. Targeting CD74 in B-cell non-Hodgkin lymphoma with the antibody-drug conjugate STRO-001. Oncotarget 14, 1–13 (2023). 10.18632/oncotarget.28341

27 Li, X. et al. Discovery of STRO-002, a Novel Homogeneous ADC Targeting Folate Receptor Alpha, for the Treatment of Ovarian and Endometrial Cancers. Mol Cancer Ther 22, 155–167 (2023). 10.1158/1535-7163.MCT-22-0322

28 Fairman, J. et al. Non-clinical immunological comparison of a Next-Generation 24-valent pneumococcal conjugate vaccine (VAX-24) using site-specific carrier protein conjugation to the current standard of care (PCV13 and PPV23). Vaccine 39, 3197–3206 (2021). 10.1016/j.vaccine.2021.03.070

29 Tuller, T., Waldman, Y. Y., Kupiec, M. & Ruppin, E. Translation efficiency is determined by both codon bias and folding energy. Proc Natl Acad Sci U S A 107, 3645–3650 (2010). 10.1073/pnas.0909910107

30 Bolanos-Garcia, V. M. & Davies, O. R. Structural analysis and classification of native proteins from E. coli commonly co-purified by immobilised metal affinity chromatography. Biochim Biophys Acta 1760, 1304–1313 (2006). 10.1016/j.bbagen.2006.03.027

31 Barta, M. L. et al. Structure of CT584 from Chlamydia trachomatis refined to 3.05 A resolution. Acta Crystallogr Sect F Struct Biol Cryst Commun 69, 1196–1201 (2013). 10.1107/S1744309113027371

32 Chu, R. S., Targoni, O. S., Krieg, A. M., Lehmann, P. V. & Harding, C. V. CpG Oligodeoxynucleotides Act as Adjuvants that Switch on T Helper 1 (Th1) Immunity. J. Exp. Med. 186, 1623–1631 (1997). 10.1084/jem.186.10.1623

33 Marques, R. F. et al. Immune System Modulation by the Adjuvants Poly (I:C) and Montanide ISA 720. Front Immunol 13, 910022 (2022). 10.3389/fimmu.2022.910022

34 Pal, S. et al. Evaluation of Four Adjuvant Combinations, IVAX-1, IVAX-2, CpG-1826+Montanide ISA 720 VG and CpG-1018+Montanide ISA 720 VG, for Safety and for Their Ability to Elicit Protective Immune Responses in Mice against a Respiratory Challenge with Chlamydia muridarum. Pathogens 12 (2023). 10.3390/pathogens12070863

35 Knudsen, N. P. et al. Different human vaccine adjuvants promote distinct antigen-independent immunological signatures tailored to different pathogens. Sci Rep 6, 19570 (2016). 10.1038/srep19570

36 Vasilakos, J. P. et al. Adjuvant activities of immune response modifier R-848: comparison with CpG ODN. Cell Immunol 204, 64–74 (2000). 10.1006/cimm.2000.1689

37 Unemo, M. et al. Sexually transmitted infections: challenges ahead. Lancet Infect Dis 17, e235–e279 (2017). 10.1016/S1473-3099(17)30310-9

38 Warfel, K. F. et al. A Low-Cost, Thermostable, Cell-Free Protein Synthesis Platform for On-Demand Production of Conjugate Vaccines. ACS Synth Biol 12, 95–107 (2023). 10.1021/acssynbio.2c00392

39 Wilding, K. M., Zhao, E. L., Earl, C. C. & Bundy, B. C. Thermostable lyoprotectant-enhanced cell-free protein synthesis for on-demand endotoxin-free therapeutic production. N Biotechnol 53, 73–80 (2019). 10.1016/j.nbt.2019.07.004

40 Stark, J. C., et al. On-demand biomanufacturing of protective conjugate vaccines Sci Adv 7, eabe9444 (2021). 10.1126/sciadv.abe9444

41 He, W. et al. Cell-free production of a functional oligomeric form of a Chlamydia major outer-membrane protein (MOMP) for vaccine development. J Biol Chem 292, 15121–15132 (2017). 10.1074/jbc.M117.784561

42 Tifrea, D. F. et al. Induction of Protection in Mice against a Chlamydia muridarum Respiratory Challenge by a Vaccine Formulated with the Major Outer Membrane Protein in Nanolipoprotein Particles. Vaccines (Basel) 9 (2021). 10.3390/vaccines9070755

43 Mueller, K. E., Plano, G. V. & Fields, K. A. New frontiers in type III secretion biology: the Chlamydia perspective. Infect Immun 82, 2–9 (2014). 10.1128/IAI.00917-13

44 Fellows, P. et al. Characterization of a Cynomolgus Macaque Model of Pneumonic Plague for Evaluation of Vaccine Efficacy. Clin Vaccine Immunol 22, 1070–1078 (2015). 10.1128/CVI.00290-15

45 Heath, D. G. et al. Protection against experimental bubonic and pneumonic plague by a recombinant capsular Fl-V antigen fusion protein vaccine. Vaccine 16, 1131–1137 (1998). 10.1016/s0264-410x(98)80110-2

46 Jneid, B. et al. Role of T3SS-1 SipD Protein in Protecting Mice against Non-typhoidal Salmonella Typhimurium. PLoS Negl Trop Dis 10, e0005207 (2016). 10.1371/journal.pntd.0005207

47 Martinez-Becerra, F. J. et al. Characterization and Protective Efficacy of Type III Secretion Proteins as a Broadly Protective Subunit Vaccine against Salmonella enterica Serotypes. Infect Immun 86 (2018). 10.1128/IAI.00473-17

48 Webster, E. et al. Immunogenicity and Protective Capacity of a Virus-like Particle Vaccine against Chlamydia trachomatis Type 3 Secretion System Tip Protein, CT584. Vaccines (Basel) 10 (2022). 10.3390/vaccines10010111

49 Ramsey, K. H., Soderberg, L. S. & Rank, R. G. Resolution of chlamydial genital infection in B-cell-deficient mice and immunity to reinfection Infect Immun 56, 1320–1325 (1988). 10.1128/iai.56.5.1320-1325.1988

50 Williams, D. M., Grubbs, B. & Schachter, J. Primary murine Chlamydia trachomatis pneumonia in B-cell-deficient mice. Infect Immun 55, 2387–2390 (1987). 10.1128/iai.55.10.2387-2390.1987

51 Putman, T. et al. ChlamBase: a curated model organism database for the Chlamydia research community. Database (Oxford) 2019 (2019). 10.1093/database/baz041

52 Pal, S., Peterson, E. M. & de la Maza, L. M. Vaccination with the Chlamydia trachomatis major outer membrane protein can elicit an immune response as protective as that resulting from inoculation with live bacteria. Infect Immun 73, 8153–8160 (2005). 10.1128/IAI.73.12.8153-8160.2005

53 Kodera, N., Sakashita, M. & Ando, T. Dynamic proportional-integral-differential controller for high-speed atomic force microscopy. Review of Scientific Instruments 77 (2006). 10.1063/1.2336113

54 Uchihashi, T., Kodera, N. & Ando, T. Guide to video recording of structure dynamics and dynamic processes of proteins by high-speed atomic force microscopy. Nat Protoc 7, 1193–1206 (2012). 10.1038/nprot.2012.047

55 Zhang, Y., Tunuguntla, R. H., Choi, P. O. & Noy, A. Real-time dynamics of carbon nanotube porins in supported lipid membranes visualized by high-speed atomic force microscopy. Philos Trans R Soc Lond B Biol Sci 372 (2017). 10.1098/rstb.2016.0226

56 Niina, T., Matsunaga, Y. & Takada, S. Rigid-body fitting to atomic force microscopy images for inferring probe shape and biomolecular structure. PLoS Comput Biol 17, e1009215 (2021). 10.1371/journal.pcbi.1009215

57 Nečas, D. & Klapetek, P. Gwyddion: an open-source software for SPM data analysis. Open Physics 10 (2012). 10.2478/s11534-011-0096-2

58 Agger, E. M. et al. Cationic liposomes formulated with synthetic mycobacterial cordfactor (CAF01): a versatile adjuvant for vaccines with different immunological requirements. PLoS One 3, e3116 (2008). 10.1371/journal.pone.0003116

59 Pal, S., Tifrea, D. F., Follmann, F., Andersen, P. & de la Maza, L. M. The cationic liposomal adjuvants CAF01 and CAF09 formulated with the major outer membrane protein elicit robust protection in mice against a Chlamydia muridarum respiratory challenge. Vaccine 35, 1705–1711 (2017). 10.1016/j.vaccine.2017.02.020

60 Pal, S., Fielder, T. J., Peterson, E. M. & de la Maza, L. M. Protection against infertility in a BALB/c mouse salpingitis model by intranasal immunization with the mouse pneumonitis biovar of Chlamydia trachomatis. Infection and Immunity 62, 3354–3362 (1994). 10.1128/iai.62.8.3354-3362.1994

